# Characterizing the performance of human leg external force control

**DOI:** 10.1101/2021.08.09.455741

**Authors:** Pawel Kudzia, Stephen N. Robinovich, J. Maxwell Donelan

## Abstract

Our legs act as our primary contact with the surrounding environment, generating external forces that enable agile motion. To be agile, the nervous system has to control both the magnitude of the force that the feet apply to the ground and the point of application of this force. The purpose of this study was to characterize the performance of the healthy human neuromechanical system in controlling the force-magnitude and position of an externally applied force. To accomplish this, we built an apparatus that immobilized participants but allowed them to exert variable but controlled external forces with a single leg onto a ground embedded force plate. We provided real-time visual feedback of either the leg force-magnitude or position that participants were exerting against the force platform and instructed participants to best match their real-time signal to prescribed target step functions. We tested target step functions of a range of sizes and quantified the responsiveness and accuracy of the control. For the control of force-magnitude and for intermediate step sizes of 0.45 bodyweights, we found a bandwidth of 1.8±0.5 Hz, a steady-state error of 2.6±0.9%, and a steady-state variability of 2.7±0.9%. We found similar control performance in terms of responsiveness and accuracy across step sizes and between force-magnitude and position control. Increases in responsiveness correlated with reductions in other measures of control performance, such as a greater magnitude of overshooting. We modelled the observed control performance and found that a second-order model was a good predictor of external leg force control. We discuss how benchmarking force control performance in young healthy humans aids in understanding differences in agility between humans, between humans and other animals, and between humans and engineered systems.

## Introduction

Agility is an important aspect of movement performance. This is true in athletics where success can be determined by how high a volleyball player jumps when blocking a hit, or how quickly a soccer player redirects their motion when taking evasive action from an oncoming defensive player. This is also true in the wild where to survive, animals must chase down their prey, evade predators, and negotiate variable terrain [1], [2]. And in robotic systems, agility will be necessary for legged robots to provide fundamental services to society that currently only humans can provide. For example, agile robots may assist in complex mountain rescues with rough and varied terrain, move payloads while avoiding obstacles in construction sites, and deliver the mail [3].

Aspects of agility are measured in several ways. In athletics, agility has been quantified by sprint speed [4], jump height [5], and time to complete an obstacle course [6], [7]. Scientists have quantified the agility of different animals and sometimes compared agility across species, by measuring maximum sprint speed [8], maximum jump height [9], and minimum turning radius [10], [11]. To accomplish any of these agility tasks—running fast, jumping high, or changing direction rapidly—requires generating well-controlled forces against the environment (external forces). In turn, the resulting reaction force from the environment accelerates our bodies. In principle, legged animals and robots can push against the ground with any part of their body, but we most commonly do this using our legs to push our feet against the ground.

To be agile, the nervous system has to control both the magnitude of the force that the feet apply to the ground, as well as the point of application of this force. The external force beneath each foot is a vector quantity. The magnitude of three orthogonal components of force must be controlled—we refer to this as force-magnitude control. As is a convention, we represent these force components as forward and backward in the horizontal plane (or anterior and posterior), side to side which is also in the horizontal plane (or medial and lateral), and upward and downward in the vertical direction. The nervous system may use a decomposition that is different from this convention. Modulating the magnitude of each component will change the acceleration of the body along that component’s direction. An accelerating sprinter selectively increases the forward component, a maximum height vertical jumper selectively increases the vertical component, and a runner changing direction may need to modulate all three components [14]. Our nervous system also needs to control the point of force application of the external force vector—we refer to the control of this point as force-position control. By selectively controlling the force-position, and components of the force-magnitude, a gymnast can change angular acceleration to initiate an aerial front flip by generating a rotational moment of force about their center of mass [15]. To control the force beneath each foot our nervous system must synergistically coordinate leg muscle activations [12], [13]. In general, the selective linear and angular acceleration of the body required for agile motion requires the rapid and accurate control of force-magnitude and force-position.

Given the importance of well controlling external force vectors to generate agile motion, we know remarkably little about how well this is accomplished by legged biological and engineered systems. One approach to characterizing the performance of a control system is to evaluate the system’s ability to rapidly and accurately change from one commanded state to a new target state, known as the step response [16]. The response to a step-change in commanded external force has been evaluated in several engineered legged systems. For example, the MIT Cheetah robot is able to step-change from zero vertical force beneath the feet to 1/3 of its bodyweight within milliseconds and does so with almost zero error [17], [18]. Similarly, a state-of-the-art foot prosthesis demonstrates millisecond-scale control of torque, allowing the prosthesis to accurately track rapidly changing commanded torques [19]. However, force control performance is not routinely measured in engineered systems, and to our knowledge, has never been quantified in humans or other legged animals.

The purpose of our study was to characterize the performance of the human leg in controlling external force-magnitude and force-position. We focused here on the control of sub-maximal forces generated by a single leg in a posture similar to how a runner actuates their motion during a run. To accomplish this purpose, we built a custom apparatus that constrained participants’ bodies from moving so that their legs maintained a constant posture while allowing them to change force-magnitude and force-position. To quantify the step response, we instructed participants to best match visually-displayed step changes in ground force magnitude by pushing more or less against a ground-mounted force plate (force-magnitude control), or step changes in ground force position by shifting the pressure beneath their foot forwards and backwards, or side to side (force-position control). We evaluated control performance by how quickly and accurately participants could match their force magnitude and force position to the commanded changes.

## Methods

### Participants

We recruited fourteen participants for the study (female: n = 4; male: n = 10; body mass: 72.2±6.1 kg; age: 27±3 years; foot length: 0.25±0.2m; mean±std). The Office of Research Ethics at Simon Fraser University approved the study and participants provided written informed consent before participating.

### Experimental Design

To characterize the performance of human leg force control, we tested the step response of participants selectively controlling leg external force. To do this, we built an apparatus that vertically and horizontally constrained participants standing over top of a ground embedded force plate (Bertec Corporation, Ohio, USA). The apparatus was rigidly attached to the ground around the force plate, but not to the force plate itself. The force plate beneath the participant’s foot sampled at a frequency of 1000 Hz and was connected to a data acquisition unit (USB-6229, National Instruments Corporation, Texas, USA), which interfaced with our computer. The constraints imposed by the testing apparatus allowed participants to exert a variable but controlled external force onto the ground by selectively pushing more or less against the ground with their leg or selectively shifting the center of pressure beneath their foot anteriorly/posteriorly (forward/backward) or medially/laterally (side-to-side). We built the testing apparatus using 1.5”x1.5” T-Slotted aluminum bars (80/20 Inc., Indiana, USA). We mounted four adjustable scissor jacks onto the apparatus to secure and immobilize the participant to the apparatus (Figure 1). Two of the scissor jacks pushed down against the shoulders, and two pushed up against the forearms. An adjustable padded bar supported the leg that was not pushing on the ground. On all points where the participant came in contact with the apparatus, we secured high-density padding to reduce discomfort.

**Figure 1.**
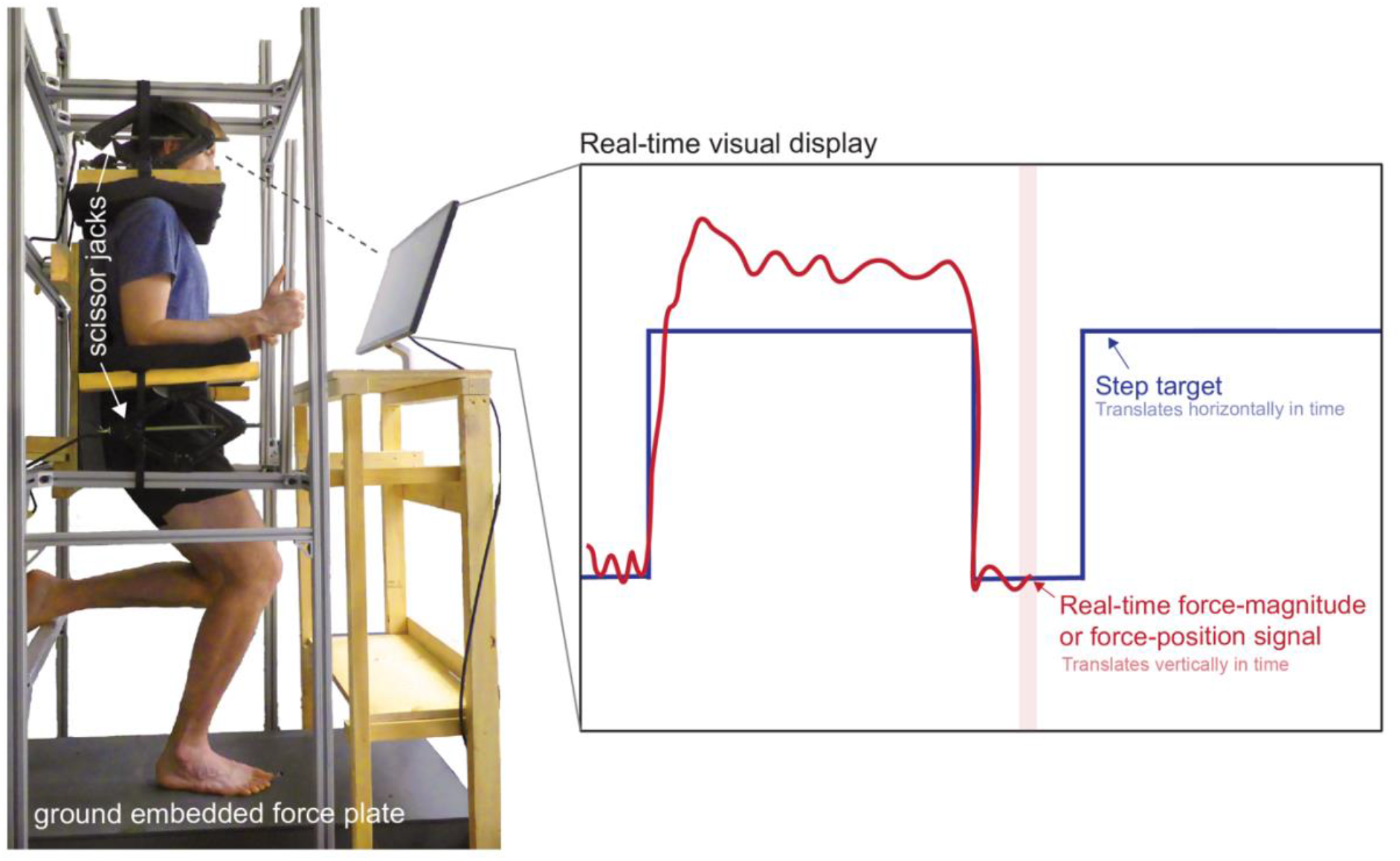
Apparatus to characterize external force control. The apparatus constrained the vertical and horizontal motion of each participants’ torso using adjustable scissor jacks that pushed down on their shoulders and pushed up against their forearms, and a stiff aluminum frame securely mounted to the ground. The participants stood in a posture resembling the stance phase of a run and could selectively push more or less or shift the pressure beneath their foot against the ground-embedded force plate. The real-time feedback displayed to the participants the real-time force-magnitude or position signal that they were exerting onto the ground as well as the target step function they were to try and best match.

To command target step changes in force-magnitude and position, we used visual feedback to allow participants to compare their target and actual force-magnitude or position. We provided real-time visual feedback using a computer monitor mounted in front of the participants (Figure 1). Our custom software (MATLAB 2019a, MathWorks) displayed both real-time feedback of either the force-magnitude or force-position signals that the participant’s foot exerted onto the force plate and of the target step function that the participant tried to best match. We filtered the raw force signal using a zero-lag low-pass fourth-order Butterworth filter with a cut-off frequency of 10 Hz. For force-magnitude control, we then normalized the real-time signal to each participant’s bodyweight and zeroed the force platform such that zero force was equivalent to one bodyweight. For the real-time force-position signal, we calculated the medial-lateral and anterior-posterior centers of pressure and filtered the raw force signal using a zero-lag low-pass fourth-order Butterworth filter with a cut-off frequency of 5 Hz. [20]. We then mathematically shifted the axes of the center of pressure on the force platform to originate beneath the participant’s foot that was in contact with the force plate. To perform this shift, we had each participant stand in a comfortable position on each leg in the rig (Figure 1). We then recorded the values of the center of pressure while participants stood on each leg and programmed a shift of the force plate axes to originate directly below the foot the participant was standing on. We programmed all the candidate real-time signals to display at the center of the screen and constrained this signal to only vertically move up and down as the participant pushed more or less against the force plate (force-magnitude control) or as they displaced their force-position anteriorly/posteriorly or medially/laterally beneath their foot (force-position control). Finally, we programmed the target step functions to slide past the real-time signal providing participants with time to view upcoming changes in the step function before they occurred (Figure 1).

We provided each participant with instructions and motivation to enable them to maximize their performance in matching their real-time force-magnitude or position signal to the targets. We instructed participants to keep their foot firmly mounted to the force platform. Our instructions to them were *“your goal in this experiment is to try and match the real-time signal to the target line that you will see on your screen. For force-magnitude control, that is pushing with force against the ground. This will mean pushing down or reducing the amount you push down, as quickly and accurately as you can to match the sliding target wave moving along the screen. For force-position control, this will mean changing the pressure beneath your foot side-to-side or forward/backward to match the sliding target wave as quickly and accurately as possible. I will notify you if it will be side-to-side or forward/backward shifting”* Due to the repetitive nature of the experiment and the possibility of both physical and mental fatigue, we provided participants with 15-second intermissions between trials (trials are explained below). During this intermission, we displayed a countdown on their screen to notify them when the next trial was beginning and a scoreboard showing them the total error incurred during each trial. We calculated this error as the total root mean square error between the empirical response and the target step function to encourage both speed and accuracy when rising and falling to new target forces and positions. We informed participants that a perfect score was 0 error. We encouraged all participants to minimize this error by best matching their real-time signal to the step targets appearing on their screen.

### Experimental Protocol

We characterized external force control over two sessions. We performed the sessions in a randomized order and on different days. In each session, we fitted participants into the experimental apparatus. When fitting participants into the apparatus, we used an analog goniometer to enforce a 15-degree knee flexion angle to approximate a posture similar to that assumed by a runner at the start of stance [21]. While maintaining this posture, we adjusted the horizontal positions of the arm and shoulder constraints such that the participant was unable to shift horizontally within the apparatus. Next, we adjusted the vertical arm and shoulder constraints by vertically shifting the scissor jacks to apply downwards pressure on the participant’s shoulders. Finally, we adjusted the height of the padded bar that supported the leg that was not pushing on the ground. In the force-magnitude session, we recorded each participant’s mass by having them stand on a force plate.

In one session, we characterized force-magnitude control. This session contained five conditions—one training and four testing. The training condition came first, and we designed it to familiarize participants with the experiment. It consisted of 12 trials, totaling six on each leg, with a single target step function size of 0.85 bodyweights. We choose to focus on a single target step function size in order to get many trials where a participant can practice all the aspects of the task including being comfortable in the apparatus and acquainting themselves with the visual display without the added complexity of changing the target size. In all trials, the target step function consisted of a square wave of ten matching size target steps, each four seconds in duration, with three seconds between steps (totaling 73 seconds per trial). The four testing conditions followed the training condition. Each of these testing conditions contained a target step function size of either 0.25, 0.45, 0.85, or 1.25. We had participants perform each of these testing conditions in random order. Testing conditions consisted of 6 trails, totaling three per leg. In all cases, participants switched the leg in contact with the ground after each trial to ameliorate the effects of fatigue.

In another session, we characterized force-position control. This session contained five conditions—one training and four testing—for anterior-posterior control and the same five conditions for medial-lateral control. In random order, we first evaluated either the conditions within anterior-posterior control or the conditions within medial-lateral control. In either case, the training condition came first, and we designed it to familiarize participants with the experiment. It consisted of 12 trials, totaling six on each leg, with a single target step function of size 2.5 centimeters anterior for anterior/posterior control trials and target step size of 1.0 centimeters for the medial/lateral control trials. Again, in all trials, the target step function consisted of a square wave of ten target steps, each four seconds in duration, with three seconds between steps. The four testing conditions followed the training condition. Each of these four conditions contained a single target step function size of either +2.5, +4.0, −1.0, −2.5 centimeters for the anterior (+)/posterior (-) control conditions or +1.0, +0.5, −0.5, −1.0 centimeters for the medial (+)/lateral (-) control conditions. Like for force-magnitude control, we had participants perform each of these testing conditions in random order. Testing conditions consisted of 6 trials, totaling three per leg and in all cases, we asked participants to swap the leg in contact with the ground after each trial.

To determine whether visual feedback slowed the participant’s ability to control external force, we eliminated visual feedback for some of the step responses in each trial. We randomly omitted the visual feedback from two of every ten steps in every trial for each condition throughout all of our experiments including the training conditions. We did this by momentarily taking away the real-time visual feedback of the force the participant is exerting onto the ground, while the target step function remains on the screen. We removed visual feedback starting from when the target steps up to when it steps back down again.

To determine how quickly force could rise when the speed of the response was prioritized over the accuracy, we had one participant perform a force-magnitude control experiment stepping up to a step target size of 0.85 bodyweights (male; body mass = 80.6 kg; foot length = 26.6 cm). Similar to the main protocol, he performed three trials with the left leg and three trials with the right leg. We instructed him that the priority was to step up as quickly to the target as they could and that it was unnecessary to match or hold the steady-state force level after stepping up to the target.

### Data Analysis

#### Step response characteristics

For each step response within each trial, we evaluated the responsiveness and accuracy in controlling external force. To accomplish this, we took the results from each trial and segmented them into ten individual step responses such that each segmentation consisted of a six-second single-step response, with one second before the step-up and 1 second after the step-down (Figure 2). For each step response, we evaluated the rise time (in units of milliseconds) as the time for the signal to go from 10% to 90% of the target step size value. We calculated the upper limit of bandwidth (in units of Hz) by dividing 0.35 by rise time [22] and assumed the lower limit was 0 Hz. The bandwidth measures the frequency range at which a target signal can change and still be accurately tracked by the controller. We quantified fall time (in units of milliseconds) as the time for the signal to go from 90% to 10% of the target step size value as the participant steps-down. In this paper, we use the term responsiveness to refer to rise time, fall time, and bandwidth—a responsive system has a short rise time, a short fall time, and a high bandwidth. Next, we quantified overshoot by taking the peak value reached by the signal, subtracting the value of the target step, dividing this by the size of the step, and multiplying by 100. In this way we expressed overshoot as the percentage that the signal exceeds the target step size value. If the signal did not surpass the target step value resulted in a negative value for the overshooting. A large overshoot can indicate under-damping in the controller—it will take longer for the system to settle and reach its target step size. We restricted our evaluation of the rise time, bandwidth, and overshoot of the response to the time interval of 0.5 seconds before the step target stepped up and up to one second after the step-up occurred. We quantified steady-state error by calculating the mean value of the signal between 2-3 seconds, subtracting the value of the target step size, and taking the absolute value. We divided this by the value of the target step size and multiplied this by 100 to express the steady-state error as a percentage of the target value. The steady-state error informs us how much error the control system has once it has reached the target size value and settled in its new state. A large error can be indicative of poor control as the system is not able to well match the target value. We quantified steady-state variability by calculating the standard deviation of the signal between 2-3 seconds, dividing this by the steady-state mean and multiplying this value by 100. The steady-state variability, expressed as a percentage of the steady-state mean, informs us how variable the system is once it has reached and settled on its new state. In this paper, we use the term accuracy to refer to overshoot, steady-state error, and steady-state variability—an accurate system has small overshoot, small steady-state error, and small steady-state variability. We evaluated the fall time between four to six seconds (Figure 2). We collectively refer to the rise time, bandwidth, overshoot, steady-state error, steady-state variability, and fall time as the step response characteristics.

**Figure 2:**
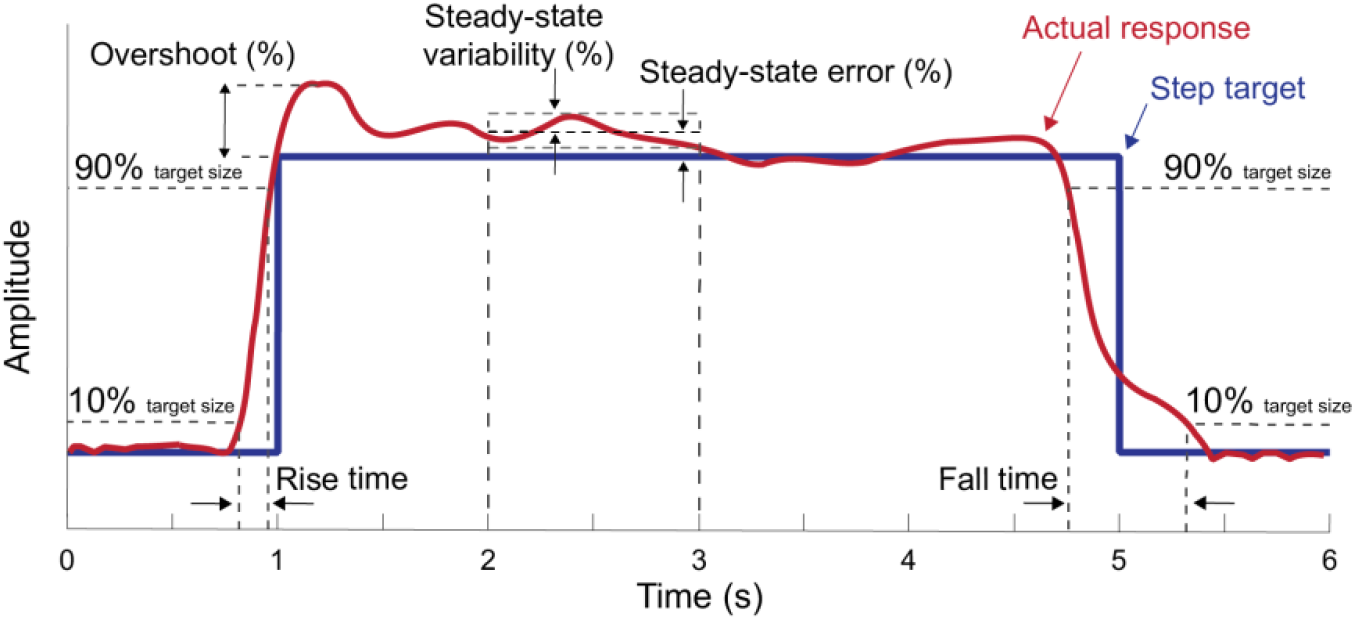
Step response characteristics. The empirical response (red line) and the step target (blue) line are shown for a single segmented target step response six seconds in length. For force-magnitude control, the step target size is in bodyweights and for force-position control, the step target size is in centimeters. The evaluated control characteristics include rise time (s), bandwidth (Hz), overshoot (%), steady-state error (%), steady-state variability (%), and fall time (s).

We used exclusion criteria to remove step responses that may poorly describe leg external force control. We used the following criteria: 1) the control response is in the opposing direction of the target step, 2) the response does not step up within 0.5 seconds after the target step function steps up, and 3) the response is already greater than 10% of the target step size value the 0.5 seconds leading up to the step-up. We removed the step responses that meet these exclusion criteria from all subsequent analyses.

#### Statistical Analysis

To estimate the fastest control performance for each participant and condition, we calculated means of step response characteristics using the five trials with the fastest rise times. We then calculated the mean and standard deviation across participants for each step response characteristic. To determine if there were differences in characteristics between different step sizes (i.e., conditions), we performed a repeated-measures analysis of variance within the force-magnitude, the medial-lateral force-position, and the anterior-posterior force-position experiments. If a difference existed, we performed pairwise comparisons between the group means to evaluate which conditions produced similar step response characteristics adjusting p-values for multiple comparisons using Bonferroni corrections [23]. To understand if there were relationships between step response characteristics within each condition, we used data from every included step response, not just the five fastest responses. We used linear mixed-effects models to identify these relationships. This approach allowed for individual y-intercepts for each participant while finding a single best-fit parameter describing the slope between the two variables. We assessed the strength of a relationship using the slope magnitude, the probability that the slope is different from zero, and the R^2^ value. In all cases, we used MATLAB’s statistical analysis toolbox and accepted p<0.05 as statistically significant.

#### System modelling

To further describe the step response characteristics of leg force control, we modelled the system dynamics and used system identification to estimate the unknown parameters. As a starting point, we modelled the relationship between the commanded step response of the input, X, and the measured step response of the output, Y, as a single dynamic process comprising a time-delayed second-order linear ordinary differential equation. We modelled this system using a second-order representation as the biological response displays overshooting, which cannot be accounted for with a simpler first-order system. The mathematical representation of the proposed model takes the form:

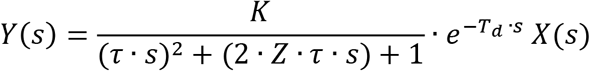

where s denotes representation in the frequency domain. K is the gain describing the steady-state value of the output Y induced by a unit change in the input X, Z is the damping constant, τ is the time constant characterizing the rate of change of Y in response to a change in X, and T_d_ is the time delay (or lead time) which corresponds to the time delay (or lead time) of when the system first begins to respond. We used the five fastest step responses for each participant and each experimental condition, used only the step-up response, and fit the model to each of those step-up responses. We normalized the magnitude of each trial to unity to allow for comparisons between different step sizes. To accomplish this normalization, we divided each step response by the size value of each step target. To fit this model to our data, we solved for the unknown model parameters by using a system identification approach. This consisted of using constrained numerical optimization (*fmincon*, MATLAB 2019a) to minimize the squared difference between the model predicted and actual empirical response (our objective function). To estimate model step response characteristics and system parameters, we followed the same statistical procedures as with the experimental step responses. That is, we determined the step response characteristics for each model fit, averaged across step responses for each participant within each condition and then averaged again across participants within a condition. We used a repeated-measures analysis of variance within the force-magnitude, the medial-lateral force-position, and the anterior-posterior force-position models and assessed the goodness-of-fit of the estimated best-fit parameters using R^2^ values.

## Results

### Step response characteristics were similar between force-magnitude and force-position control

When controlling the force-magnitude of leg external force, participants were quickest to reach the smallest target step size. At the smallest step size of 0.25 bodyweights, participants stepped up to the target at a rise time of 178±69 ms and stepped down with a fall time of 399±122 ms (Table 1, and Figure 3,4). This rise time equated to a bandwidth of 2.2±0.6 Hz. Once participants did reach the commanded target step size, they did so by overshooting it by 31.6±15.8%. Upon settling at the target, participants did so with a steady-state error of 3.9±0.4% and a steady-state variability of 4.4±1.5%. As the force-magnitude target step size increased, participants took longer to reach the target resulting in decreases in bandwidth (Table 1). Participants also overshot the target less and were, in general, more accurate and less variable during steady-state (Table 1). Fall times were always longer than rise times (p<0.001), and we observed no changes in fall times with increasing target step size.

**Table 1:** Average control characteristics for force-magnitude and force-position control

| Vertical |  | Force-magnitude control |  |  |  |  |
| --- | --- | --- | --- | --- | --- | --- |
| Target step size<br>( bodyweight) | Rise time (ms) | Fall time (ms) | Bandwidth (Hz) | Overshoot (%) | SSE (%) | SSV (%) |
| 0.25 | 177.6 ± 69.3 | 399.2 ± 121.9 | 2.2 ± 0.6 | 25.3 ± 12.7 | 3.2 ± 1.0 | 3.5 ± 1.2 |
| 0.45 | 204.7 ± 63.5 | 373.8 ± 99.8 | 1.8 ± 0.5 | 17.1 ± 9.5 | 2.6 ± 0.9 | 2.7 ± 0.9 |
| 0.85 | 242.9 ± 92.3 | 389.8 ± 112.1 | 1.6 ± 0.5 | 9.8 ± 5.6 | 2.4 ± 1.3 | 2.1 ± 0.4 |
| 1.25 | 266.0 ± 90.5 | 405.5 ± 99.0 | 1.4 ± 0.4 | 7.4 ± 3.3 | 2.6 ± 1.3 | 2.5 ± 0.7 |
| p-value | <0.001 | 0.8 | <0.001 | <0.001 | 0.26 | <0.001 |

| Anterior (+) Posterior (-) |  | Force-position control |  |  |  |  |
| --- | --- | --- | --- | --- | --- | --- |
| Target step size<br>(cm) | Rise time (ms) | Fall time (ms) | Bandwidth (Hz) | Overshoot (%) | SSE (%) | SSV (%) |
| -1.0 | 181.1 ± 29.0 | 377.3 ± 155.1 | 2.0 ± 0.3 | 45.0 ± 17.8 | 13.9 ± 4.7 | 8.6 ± 2.8 |
| -2.5 | 225.9 ± 47.6 | 417.6 ± 111.3 | 1.6 ± 0.3 | 19.4 ± 9.2 | 6.2 ± 2.1 | 3.8 ± 2.3 |
| 2.5 | 217.4 ± 42.6 | 411.8 ± 128.0 | 1.7 ± 0.3 | 20.8 ± 10.9 | 6.7 ± 3.3 | 5.0 ± 2.0 |
| 4.0 | 244.2 ± 46.8 | 423.3 ± 100.7 | 1.5 ± 0.3 | 15.8 ± 11.1 | 4.9 ± 2.3 | 4.8 ± 2.4 |
| p-value | <0.001 | 0.76 | <0.001 | <0.001 | <0.001 | <0.001 |

| Medial (+) / Lateral (-) |  | Force-position control |  |  |  |  |
| --- | --- | --- | --- | --- | --- | --- |
| Target step size<br>(cm) | Rise time (ms) | Fall time (ms) | Bandwidth (Hz) | Overshoot (%) | SSE (%) | SSV (%) |
| 0.5 | 200.4 ± 59.6 | 377.5 ± 103.1 | 1.9 ± 0.4 | 36.0 ± 14.1 | 9.1 ± 3.4 | 6.1 ± 2.4 |
| -0.5 | 192.4 ± 39.0 | 312.2 ± 140.9 | 1.9 ± 0.3 | 32.1 ± 16.3 | 10.0 ± 4.2 | 5.5 ± 2.0 |
| -1 | 229.4 ± 37.0 | 349.0 ± 115.4 | 1.6 ± 0.2 | 15.6 ± 6.2 | 5.5 ± 2.2 | 3.3 ± 1.1 |
| 1 | 230.6 ± 45.7 | 382.1 ± 114.5 | 1.6 ± 0.3 | 19.0 ± 12.8 | 5.9 ± 3.7 | 4.2 ± 2.5 |
| p-value | <0.001 | 0.3 | <0.001 | <0.001 | <0.001 | 0.001 |

Note: Steady-state error (SEE), Steady-state variability (SSV)

**Figure 3:**
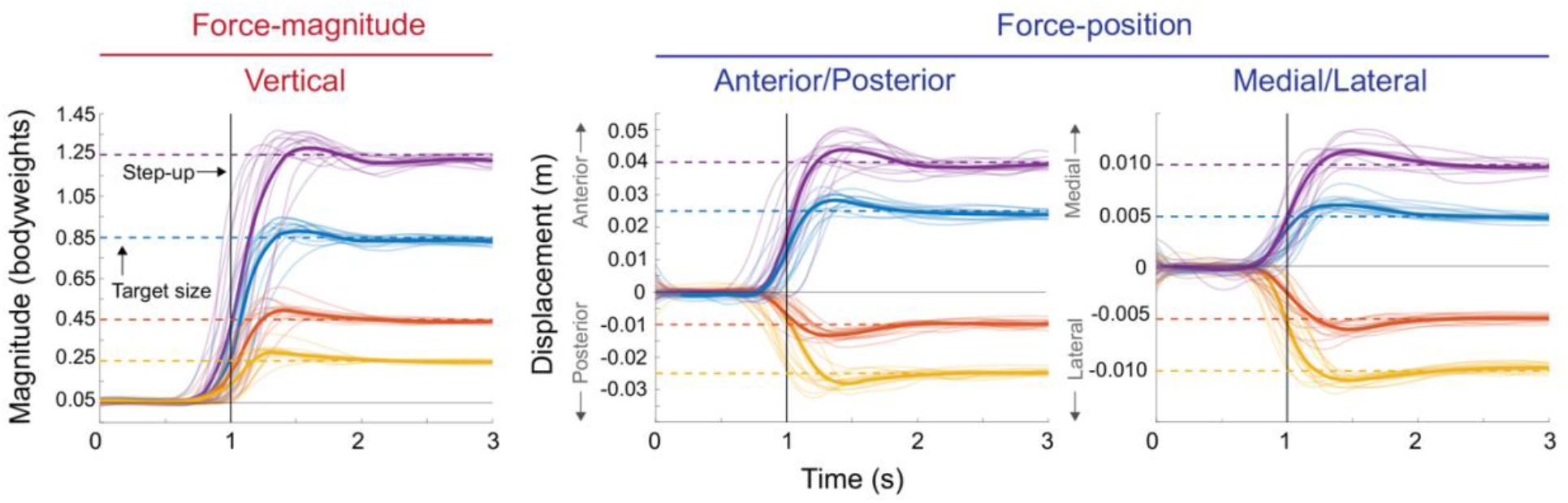
The average step-up response from each participant at each target step size (color lines) and the average of the average response (bolded color lines). The vertical line represents when the target step function steps up from its base value of 0.05 bodyweights for force-magnitude control or 0 cm for force-position control to a new target step size (dashed lines). Participants could anticipate when the step target would step up to a new target size therefore the response could precede the visual step change. Here the first second before the step-up and the two seconds after are shown. Within this time frame, we evaluated the rise time, bandwidth, overshoot, steady-state error, and steady-state variability.

**Figure 4:**
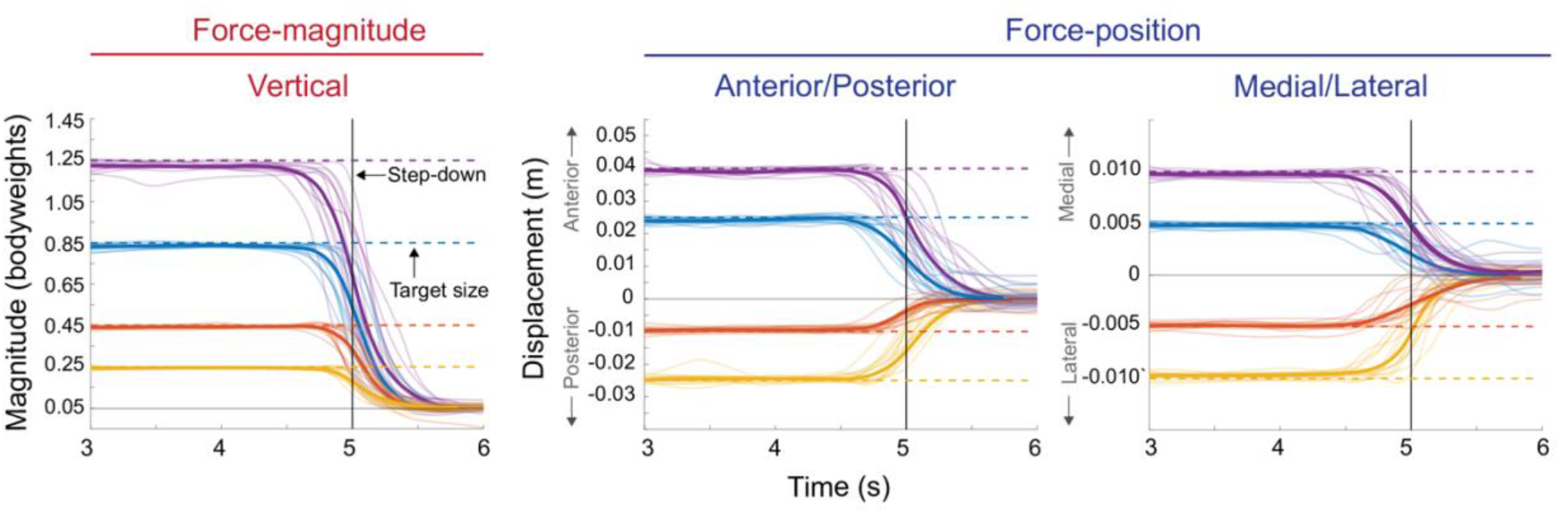
The average step-down response for each participant at each target step size (color lines) and the average of the average response (bolded colored lines). The vertical line represents when the step function steps down from the current step target size (dashed lines) to 0.05 bodyweights for force-magnitude control or 0 cm for force-position control (base values). Here the two seconds before the step-down are shown and the one second after the signal returns to its base value. Within this timeframe, we evaluated the fall times as participants stepped down from the step target size back down to the base value.

The characteristics of force-position control were comparable to force-magnitude control. When participants shifted their force-position anteriorly or posteriorly, or when they shifted their force-position medially or laterally, they did so with a rise time and bandwidth that was similar to when controlling for force-magnitude. Fall times were similar to those found for force-magnitude control and also did not change with changes in target step size for both anterior-posterior and medial-lateral control (Table 1, Figure 3,4). As in force-magnitude control, participants overshoot the target less and were in general, more accurate and less variable during steady-state with increases in target step size in force-position control (Table 1).

### A faster response correlated with reductions in other measures of control performance

On average, trials with higher bandwidth had a larger overshoot (Figure 5). For force-magnitude control, the relationship between the bandwidth of the response and overshooting the target was significant for all conditions (p<0.001). The strongest relationship had a linear fit with an *R*^2^ of 0.51 for target step sizes of 0.45 bodyweights, having a slope of +15.4% overshoot/Hz; the weakest relationship had a linear fit with an *R*^2^ of 0.36 for target step sizes of 1.25 bodyweights, and a slope of +8.5% overshooting/Hz. For force-position control, this relationship was also significant for all conditions (p<0.001) with an average fit *R*^2^ of 0.34±0.1, and an average slope of +17.2±3.6% overshooting/Hz (representative conditions are shown in Figure 5). With increasing bandwidth, participants tended to be less accurate and more variable as demonstrated by increased steady-state error and variability, but these relationships were weak and not significant for most of the conditions. Finally, there were significant correlations between increases in steady-state variability and increases in steady-state error (p<0.001) for all the conditions for both force-magnitude and force-position control, but the fits were weak with *R*^2^ values at best 0.34 and at worst 0.05. We found no other significant correlations between other measures of control performance.

**Figure 5:**
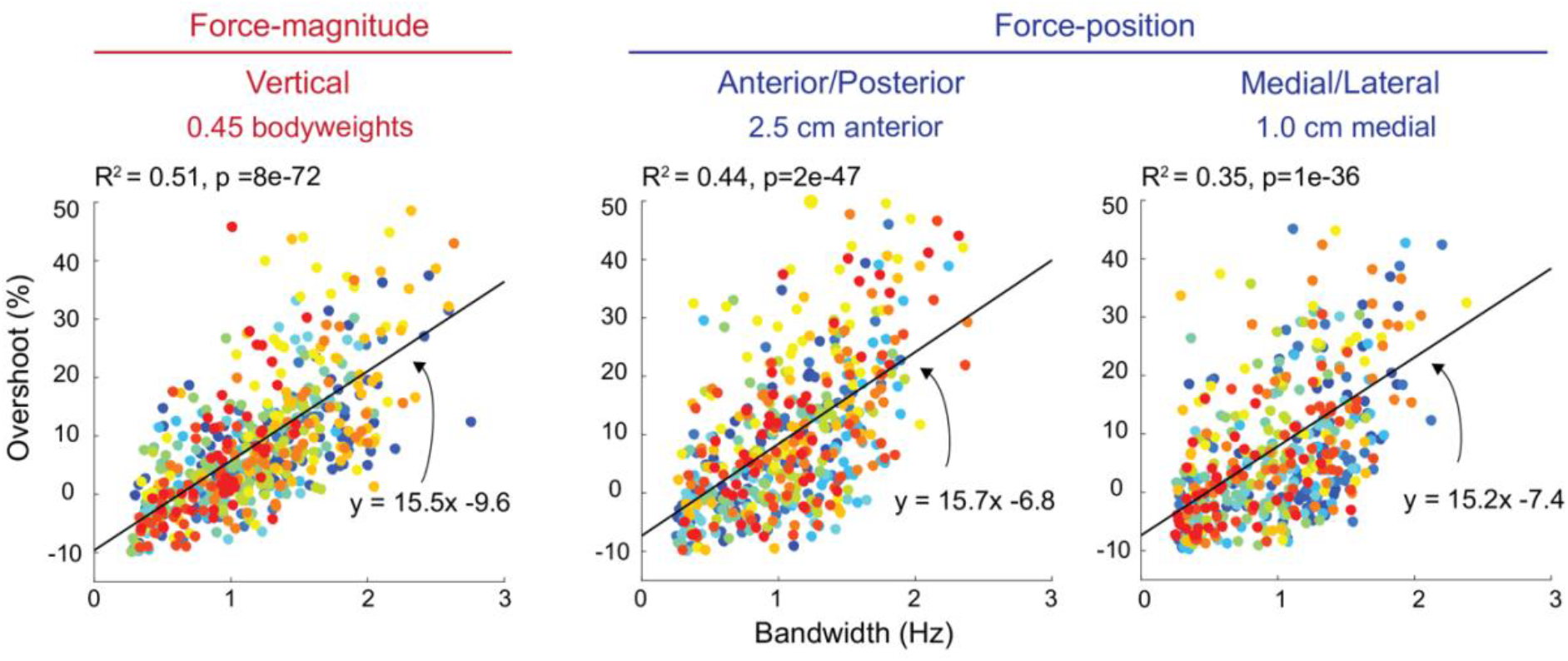
Representative findings from force-magnitude control (0.45 bodyweights), and force-position control (2.5 cm anterior and 1.0 cm medial). We fit a linear mixed-effects model to the data and plot the resultant linear fit as the black line. Each data point corresponds to the results from a single-step response while each color corresponds to a different participant. The R^2^ value for the fit is shown alongside the p-value, and equation of the line.

### Leg external force control as a second-order control system

A second-order model well described the step-up control characteristics found for the control of leg external forces (Figure 6). Best-fit R^2^ values for all step target sizes in both force-magnitude and position control were above 0.85 (Table 2). We chose the second-order model because the overshooting observed in the data was not a feature that could be described using a first-order model, and a third-order model was unnecessary given how well the second-order model described the observed behavior. The second-order controller for controlling leg external force can be well approximated using the following control parameters: K (process gain) = 1, τ (time constant) = 100 ms, Z (damping constant) = 0.5, and T_d_ (time delay or lead time) = −100ms. A controller with these parameters has a rise time of 165 ms, a bandwidth of 2.0 Hz, overshoots by 16.3%, a settling time of 1.7s, and no steady-state error or steady-state variability.

**Figure 6:**
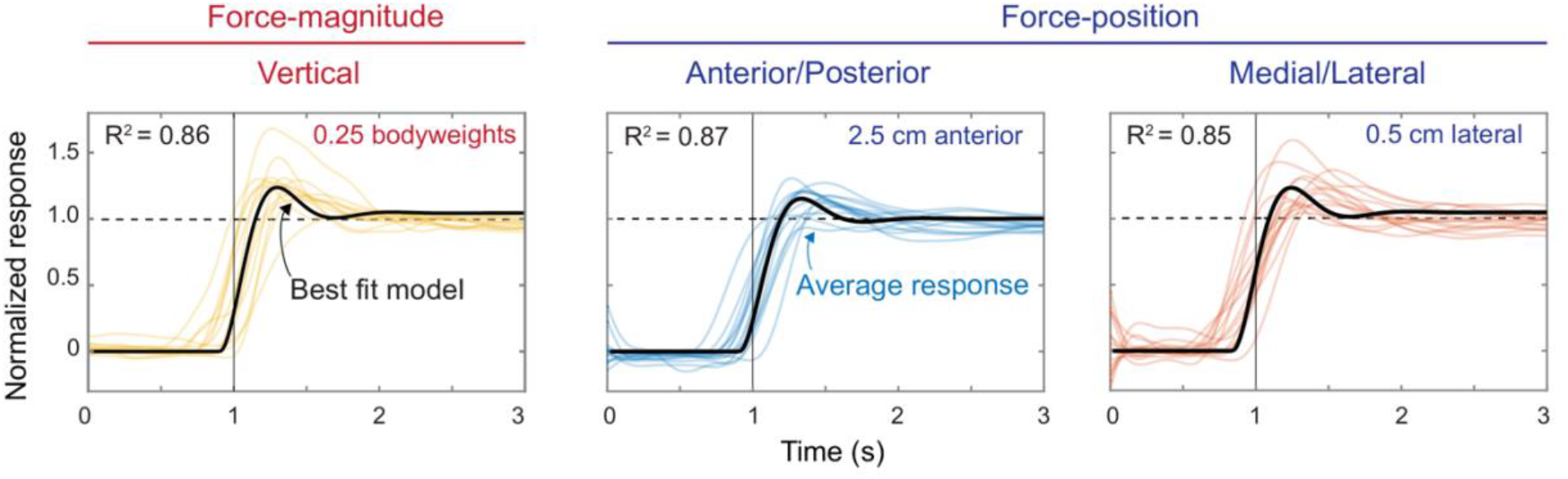
Representative modelling results for select force-magnitude and force-position target step size conditions. We plotted the second-order model using the average of the best-fitted parameters (black line) against the average of each participant’s response to each step target magnitude (colored lines). The vertical line represents when the step function steps up from zero to the normalized target value of one. The horizontal dashed line is the target step size value of 1. The R^2^ value shown is the average fit of the model for that condition.

**Table 2:**
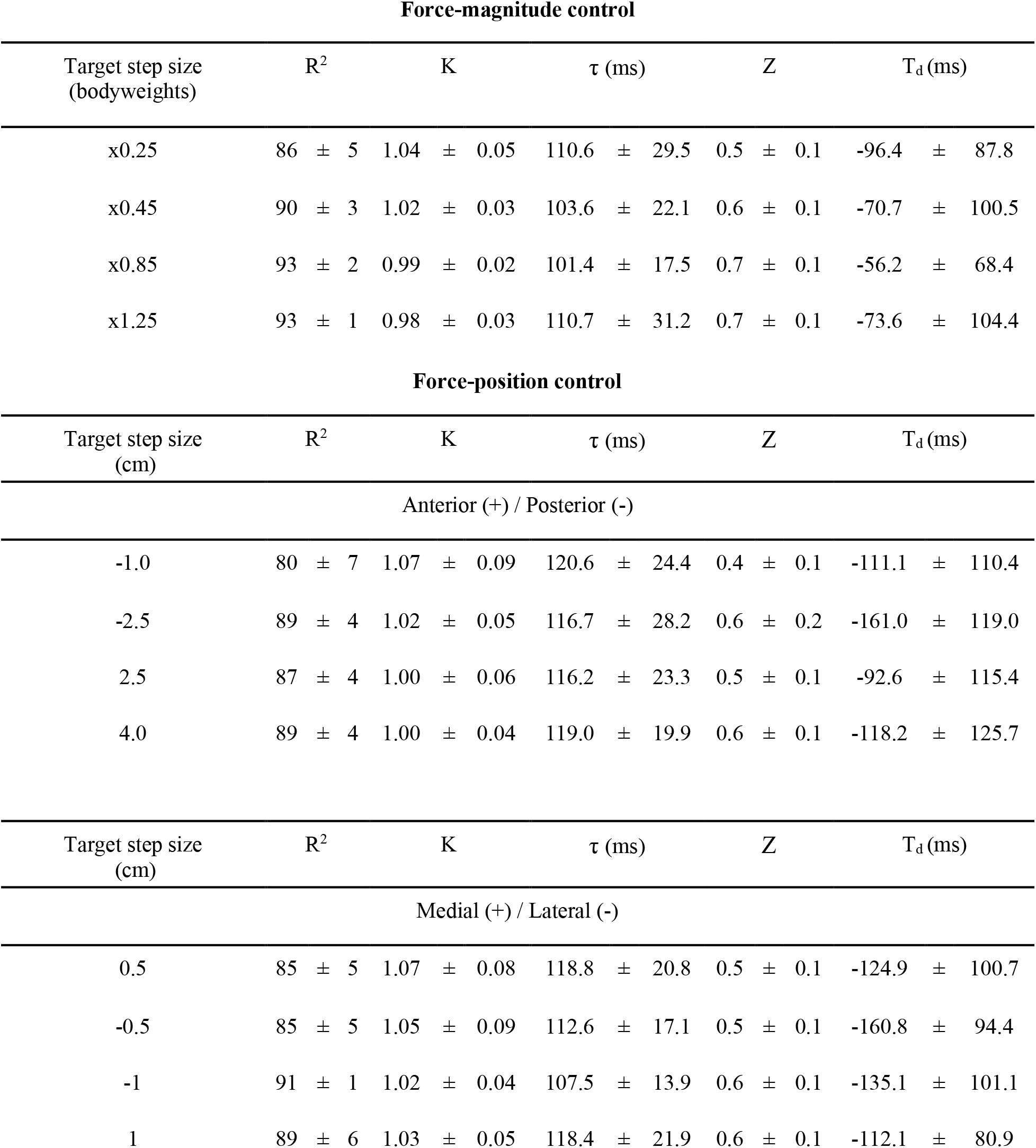
Average best fitting system modelling parameters describing leg force control as a second-order system. The R^2^ represents the fit, K is the gain used to determine error (1 being no error in the system), τ is the time constant characterizing the rate of change of how quick the system changes state, Z is the damping constant where a value of < 1 indicates an underdamped system and T_d_ is the time delay (or lead time) which is the time delay of when the system first begins to respond (a negative value indicates the system is predicting and responding before the step target steps up).

### Practice trials were effective at training participants to perform the task

To determine whether the amount of training was sufficient for participants to reach state-state, we compared the performance of their last 15 practice steps to the 15 steps immediately prior. We found no significant improvements in the rise times when comparing these two sets of trials in either the force-magnitude task trials (average difference of 17±275 ms, p=0.40) or the force-position control in the anterior/posterior direction (average difference of 2±392 ms, p=0.93) and the medial/lateral direction (45±377 ms, p=0.09). The allotted practice appears sufficient for participants to have reached a steady-state level of performance.

### Removing visual feedback resulted in specific and modest increases in performance

To test whether the task’s reliance on visual feedback affected force control performance, our protocol periodically eliminated the step that was displayed on the screen, removing the participant’s ability to use visual feedback. For force-magnitude control, we found that participants were indeed faster without visual feedback. When comparing the rise times that participants stepped up to the step target without visual feedback to that with visual feedback participants were on average 19% faster (p=0.02) for a step target size of 0.25 bodyweights, 15% faster (p=0.04) for a step target size of 0.45 bodyweights, 13% faster (p=0.02) for a step target size of 0.85 bodyweights, and 10% faster (p=0.13) for a step target size of 1.25 bodyweights. This increase in speed with the removal of visual feedback did not carry over to force-position control—we found that participants were, if anything, slower at controlling external force-position for either the anterior/posterior and medial/lateral control at all target step sizes.

### Prioritizing speed over accuracy increased the responsiveness of force control but reduced accuracy

To determine how quickly force could rise when the speed of the response was prioritized over the accuracy, one participant performed a force-magnitude control experiment stepping up to a target step size as fast as possible under instructions that removed the need to hold an accurate steady-state force level. Under these conditions, this participant had a rise time of 121±28 ms, a bandwidth of 3.0±0.49Hz and overshot the target by 50.7±19.5%. This rise time was ~100% faster than when completing it under the previous instructions (rise time of 243±92 ms) where we emphasized the need for both speed and accuracy. We observed a more than five-fold increase in overshooting the target value over completing this task under previous instructions (overshot by 9.8±5.6%) indicating a reduction in the accuracy.

## Discussion

Here we characterized people’s ability to control leg external forces. We found that people could control the magnitude of the force that they applied to the ground, and the position of that force, by responding to and closely matching commanded step changes. For both force-magnitude and position control, we observed similar control performance in terms of responsiveness and accuracy. We also observed that increases in responsiveness correlated with reductions in other measures of control performance, such as a greater magnitude of overshooting. We modelled the observed control performance and found that a second-order model was a good predictor of external leg force control.

Our study has several limitations. One limitation is that compliance—both within our participants’ bodies and within the apparatus itself—may have slowed the change in measured force-magnitude and position. To address this, we designed the apparatus using high-density padding at the contact points and adjustable scissor jacks that allow us to tightly secure each participant within the apparatus. We also reduced the possible bending of the apparatus itself by using stiff aluminum for the frame and bolting the rig securely to our laboratory floor. While it is impossible to entirely remove the effect of compliance, we aimed for our design to render the effect small.

A second limitation is that reliance on visual feedback may have slowed measured control performance. Visual feedback loops have longer loop delays when compared to spinal reflex loops, and spinal reflexes are perhaps sufficient for the neural control of leg force during tasks like running [24], [25]. Consequently, in part of our experimental design, we tested to see if control without visual feedback would be significantly faster. Indeed, we found that trials that did not rely on visual feedback were modestly faster suggesting that our measurements are a reasonable but slightly slow approximation of control performance when controlling leg external forces with faster spinal reflexes.

A third limitation is that our experiment may not adequately capture the control approach our nervous system is using when having to rapidly control leg external forces. In support of this possibility, sprinters run with a step frequency above 4 Hz [26] whereas our experimental results suggest that force control bandwidth should not exceed about 2 Hz. One mechanism for faster control is to not rely on visual or even spinal reflex loops but instead for the nervous system to estimate the motor commands in advance and execute them in a feedforward manner. To test whether force control could indeed be performed more rapidly when the need for feedback is removed, we instructed a single participant to prioritize leg force control responsiveness over accuracy when following commanded step changes in force. We found an increase in responsiveness of ~100% but a five-fold decrease in accuracy suggesting that controlling external forces in a feedforward manner can be considerably faster than when relying on feedback control but is much less accurate. As feedback is essential for accurate performance in the face of uncertainty, this observed increase in performance is likely an overestimate of the combined feedforward and feedback approach our control system likely employs under most situations [27],[28].

Sensorimotor delays may limit responsiveness of leg external force control. An inherent property of biological control systems is that there are neural delays associated with sensing and transmitting neural information, and muscular delays associated with force generation. Neural delays arise from the times required to sense a stimulus, transmit neural signals along the length of nerves, and to cross synapses [29], [30]. Muscular delays arise from the times required for conducting action potentials along muscle fibers, generating muscle force, and muscle shortening to stretch tendons that act in series [29], [30]. Using scaling relationships from other terrestrial mammals, we estimate the human delays to be approximately 1 ms for sensing, 20ms for sensory nerve conduction delay, 1 ms for crossing a single synapse in the spinal cord, 20 ms for motor nerve conduction delay, 1ms for crossing the neuromuscular junction, 10 ms for the electromechanical delay, and 40 ms for the force generation delay [29], [30]. This equates to an estimate of total human sensorimotor delay of ~90 ms, which corresponds well to estimates derived from human measurements [31]–[33]. Both feedback and feedforward control of leg external force must contend with the presence of sensorimotor delays which may set limits to responsiveness. In the feedback control approach, if the delays grow too large relative to the period of the movement, this could destabilize the system as the control signals become outdated, and the motor commands generated are no longer appropriate. To remain stable, a delayed feedback controller must use low gains resulting in low generated forces and a responsiveness that is potentially much slower than the delay itself. This may explain why a 90 ms sensorimotor delay results in controlled step changes in force that we measured here, which have a minimum period of ~500ms (2Hz bandwidth). Feedforward control of external force can generate faster responses because delays do not affect its stability. But the rate at which our muscle actuators can change between different force levels is still slow thus also limiting the bandwidth of feedforward control.

Sensorimotor noise may limit leg external force control accuracy. Noise is the random or unpredictable disturbances to a signal which obscure and interfere with the transfer of information [34]. Like delays, there are many sources of noise in our neuromechanical system. It has contributions from both neural and muscular sources. Neural sources of noise arise during the processes associated with determining movement-appropriate motor commands. These include but are not limited to noise in the uncertainty of sensory feedback, and probabilistic behavior of both cellular and synaptic signal transduction in both the peripheral and central neural systems [34], [35]. Muscular sources of noise arise during the transformation of motor commands to the contraction of muscles. Variability in the temporal structure of the motor commands and in the recruitment properties of motor units result in muscular noise [36]. Even when motor commands are perfectly timed, there is natural variability in the muscle forces arising from the stochastic nature of the contraction of sarcomeres within muscle tissue [34]. The net effect of this sensorimotor noise is that for a fixed command to the muscles, there is variability in force output. This sensorimotor noise is believed to limit performance in various human motor control tasks. For example, in goal-directed arm movements, variability in the endpoint position has shown to be related to the sensorimotor noise in the execution of the movement [35]. In isometric force production of the fingers, increases in external force magnitude show increases in force variability [36]. As with these other tasks, sensorimotor noise may limit the accuracy for leg external force control to the levels we measure here.

Measurements of force control performance may help in understanding differences in agility between humans and engineered systems, between humans and other animals, and between humans and other humans. The effective control of leg external forces in humans appears to be much slower and less accurate when compared to some legged engineered systems [3]. For example, the legs of the MIT cheetah robot have a force-magnitude bandwidth of ~100 Hz—about 50x faster than what was found in our experiment— and nearly no steady-state error or steady-state variability [17], [37]. Wearable devices such as foot prostheses and leg exoskeletons demonstrate millisecond-scale control, enabling for accurate and rapid control of force [19], [38]. Yet the agility of humans continues to be greater than that of state-of-the-art legged robots suggesting that human agility is achieved not through a greater control of leg forces, but by a greater understanding of what forces to apply. This may not hold true for comparing humans to other agile animals, such as gazelles or cheetahs, which may understand what forces to apply to the ground as well or better than humans. And in some cases, better control of external forces may be responsible for the greater agility of some animals over other less agile animals. The same may hold true for more agile athletes over less agile athletes, and for the changes in agility that come with fatigue, injury, disease, and age. Our work here benchmarks force control performance in young healthy humans to better enable these future comparisons.

## Acknowledgements

We thank all participants for volunteering their time in our study and to the SFU Locomotion Lab members and Professor James Wakeling for helpful comments and suggestions. We thank Dr. Heather More with assistance in editing the figures.

## Author Contributions

P.K, S.R, and J.M.D designed the study. P.K collected, processed, and analyzed the data. P.K. prepared all figures. P.K and J.M.D wrote the manuscript and P.K, S.R and J.M.D edited manuscript.

## Competing interests

The authors declare no competing or financial interests.

## Funding

This work was supported by the Natural Sciences and Engineering Research Council of Canada Discovery Grant (RGPIN-2020-04638) to J.M.D and the NSERC PGS Doctoral Scholarship and the SFU Graduate scholarship to P.K.

## Data Availability

The datasets generated during and/or analysed during the current study are available from the corresponding author on reasonable request.

## Addition Information

**Correspondence** and requests for any materials should be addressed to J.M.D.

## Disclosures

None

## Notes

### Competing Interest Statement

The authors have declared no competing interest.

